# Two microRNAs are sufficient for embryogenesis in *C. elegans*

**DOI:** 10.1101/2020.06.28.176024

**Authors:** Philipp J. Dexheimer, Jingkui Wang, Luisa Cochella

## Abstract

The Microprocessor, composed of Drosha and Pasha/DGCR8, is necessary for the biogenesis of canonical microRNAs (miRNAs) and essential for animal embryogenesis. However, the cause for this requirement is largely unknown: the Microprocessor may be required to produce one or few essential miRNAs, or many individually non-essential miRNAs. Additionally, Drosha and Pasha/ DGCR8 may be required for processing non-miRNA substrates. To distinguish between these possibilities, we developed a system in *C. elegans* to stringently deplete embryos of Microprocessor activity. Microprocessor-depleted embryos fail to undergo morphogenesis or form organs. We show that this early embryonic arrest is rescued by the addition of two miRNAs from the miR-35 and miR-51 families, resulting in morphologically normal larvae. Thus, just two canonical miRNAs are sufficient for morphogenesis and organogenesis, and the processing of these miRNAs accounts for the essential requirement for Drosha and Pasha/DGCR8 during *C. elegans* embryonic development.

---

MicroRNAs (miRNAs) are a class of post-transcriptional repressors with essential roles in animal development (*1*). Loss of Drosha, Pasha/DGCR8 or Dicer, required for miRNA biogenesis, or the Argonaute effector proteins, required for target mRNA silencing, results in early developmental lethality in every animal species studied (*2*). Animals often express hundreds of miRNAs, and it remains unclear what fraction of those are essential, individually or in combination. In zebrafish, the abundant miR-430 accounts for a large fraction of the embryonic defects observed upon loss of Dicer, most prominently abnormal brain formation (*3*), acting at least in part by clearing maternal mRNAs (*4*). However, miR-430 is only present in vertebrates, and in zebrafish in particular it forms an unusual cluster with >50 genomic copies (*5*). In other animals, the reasons for the Microprocessor requirement remain unclear.

To investigate the role of the Microprocessor during embryogenesis of the nematode *C. elegans*, we developed a method for stringent removal of Drosha and Pasha/DGCR8, and thus miRNAs, in embryos. Microprocessor depletion in embryos is challenging, as homozygous mutant embryos derived from heterozygous Drosha (*drsh-1*) or Pasha/DGCR8 (*pash-1*) mutant mothers carry enough maternal RNA and/or protein to develop into adults, albeit sterile (*6*, *7*). We developed a conditional approach using the *Arabidopsis* Auxin Inducible Degradation (AID) system (*8*). To this end, we tagged endogenous Drosha and Pasha/DGCR8 with the TIR1 ubiquitin ligase-recognition peptide, and expressed TIR1 in both germline and soma to clear maternal and zygotic tagged proteins **(Fig. 1A, S1A)**. To further reduce Microprocessor levels, we simultaneously triggered systemic RNAi against *pash-1* **(Fig. S1B)**. To ensure elimination of maternal miRNAs, mothers were grown on auxin and RNAi-eliciting bacteria for 20 hours before harvesting just-fertilized embryos (2-cell stage). Whereas depletion of either PASH-1 or DRSH-1 alone was sufficient to induce penetrant embryonic lethality **(Fig. S1C)**, simultaneous removal of both proteins as well as *pash-1* mRNA resulted in the earliest, most homogeneous arrest phenotype **(Fig. S1D, E)**. We refer to this combined AID and RNAi manipulation as RNAiD for simplicity. This system enabled the examination of *C. elegans* embryos that contained only trace amounts of miRNAs **(Fig. 1A, B)**; 100% of these embryos arrest uniformly at the end of gastrulation/onset of morphogenesis, lacking distinguishable internal structure, and eventually die **(Fig. 1A, C, S1D, E)**.

**Fig. 1.**
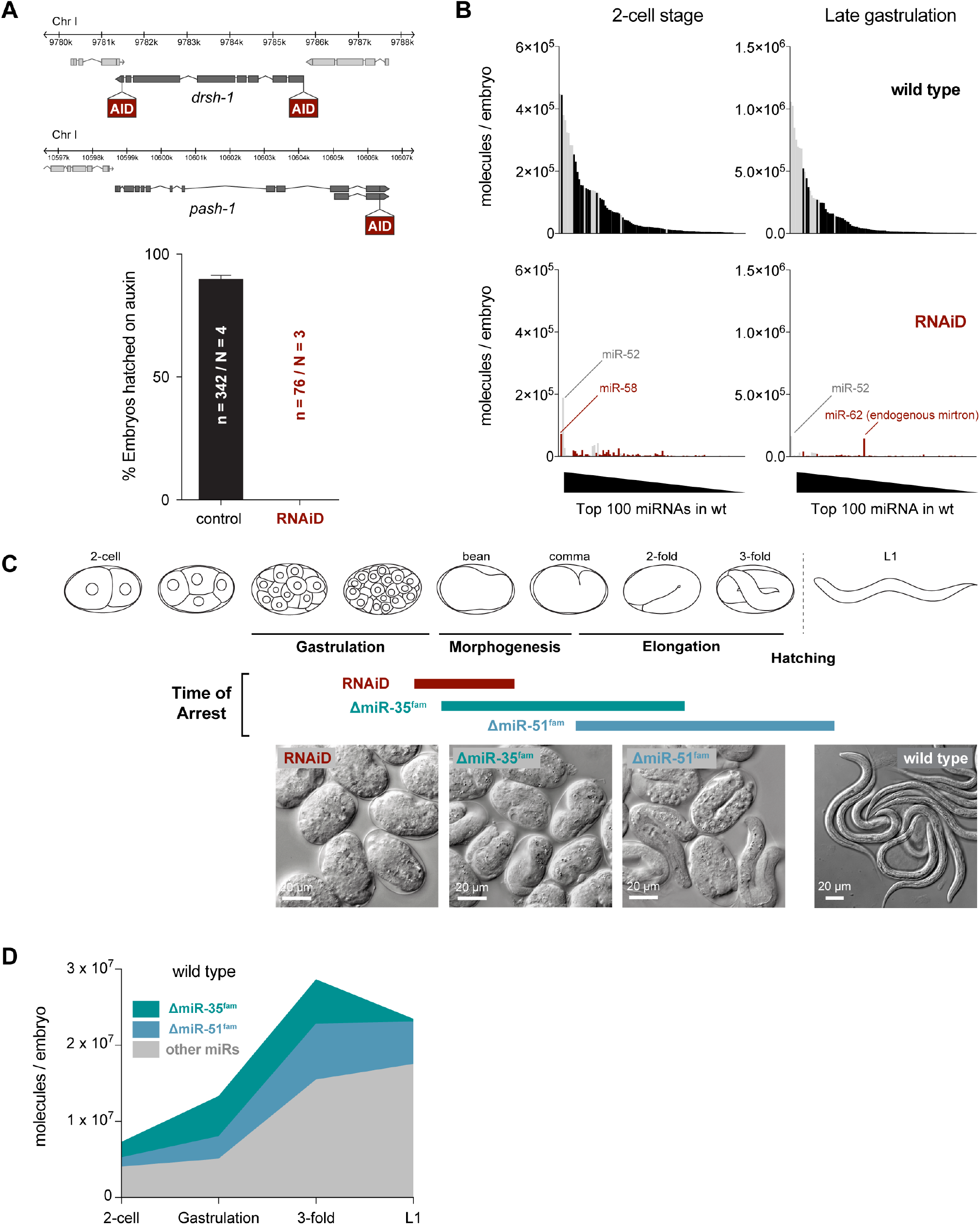
**A.** Schematic of the genetic setup to deplete the Microprocessor via the Auxin-inducible degron (AID) system; and hatching rate of wild-type or Microprocessor-depleted embryos by AID and RNAi against *pash-1* (RNAiD). n = number of embryos scored, N = number of independent experiments. Error bars represent the standard error of the proportion. **B.** Absolute miRNA abundance in wt and RNAiD embryos at the 2-cell stage and at the end of gastrulation as determined by quantitative small RNA-sequencing using a series spike-in oligos (see methods). The 100 most abundant miRNAs in wt embryos at each stage are shown. Remaining miRNAs of the miR-35 and -51 families are in gray. **C.** Schematic of the predominant time of arrest and representative images of embryos of the indicated genotypes. **D.** Absolute miRNA abundance in developing wt embryos measured by small RNA-seq, highlighting the proportion of miR-35 and miR-51 family members.

To conditionally deplete the Microprocessor by independent means, we used *pash-1(mj100)*, a temperature-sensitive allele of Pasha/DGCR8 (*9*). Shifting mothers to the restrictive temperature 7 hours before harvesting 2-cell embryos (the maximum duration possible before the mothers visibly deteriorate) also resulted in fully penetrant embryonic lethality **(Fig. S1F)**. However, the timing of arrest was variable **(Fig. S1D, E)**, consistent with the observation that at the 2-cell stage an average of 36% of maternal miRNA content remained **(Fig. S1G)**. Thus, while the *pash-1(ts)* allele is useful for removing miRNAs from mid-gastrulation onwards, RNAiD results in stringent miRNA depletion up to late gastrulation (just before embryos arrest).

In *C. elegans,* no individual miRNA is known to be essential for embryogenesis, but two families with multiple, redundant members each, miR-35^fam^ and miR-51^fam^, are necessary at two distinct timepoints (*10*–*12*). Deletion of all eight members of the miR-35^fam^ causes arrest shortly after completing gastrulation during early morphogenesis, while animals lacking the six miR-51^fam^ members have later elongation and organogenesis defects, and arrest as late embryos or malformed larva **(Fig. 1C, S1D, E)**. These two miRNA families constitute ~50% of the embryo miRNA contents **(Fig. 1D)** and they are broadly, if not ubiquitously expressed during embryogenesis (*11*, *12*). The remaining >100 miRNAs are individually much less abundant and tend to be expressed in a highly cell-type-specific manner (*2*, *13*, *14*).

The effect of RNAiD was more severe than the arrest phenotype of animals lacking the early-acting miR-35^fam^ **(Fig. 1C, S1D, E)**, raising the possibility that other miRNAs could be necessary for earlier stages of embryogenesis. Alternatively, non-miRNA related activities of Drosha and Pasha/DGCR8 could be responsible for the more severe phenotype upon RNAiD. Finally, it is possible that the combined loss of miR-35^fam^ and miR-51^fam^ accounts for the observed defects in embryo-genesis in the absence of the Microprocessor.

To distinguish between these possibilities, we asked whether miR-35^fam^ and miR-51^fam^ are sufficient for embryogenesis in the absence of all other miRNAs. As every individual member of the miR-35 or miR-51 family can rescue the complete deletion of each respective family (*11*), we reasoned we could re-introduce one member of each family into Microprocessor-depleted animals to test their sufficiency. Previous approaches to introduce individual miRNAs in a Microprocessor-independent manner relied on injection of processed RNA duplex (*3*). Instead, we developed a transgenic strategy for Microprocessor-independent miRNA delivery, which recapitulates the spatio-temporal specificity of expression of the re-introduced miRNAs. Specifically, we designed miR-35^fam^ and miR-51^fam^ mirtrons, which are processed by the spliceosome and subsequently cleaved by Dicer to produce mature miRNAs in a Microprocessor-independent manner **(Fig. 2A, Fig. S2)** (*15*, *16*). Mirtron-35 and mirtron-51 (mirt-35/51) were expressed under the respective miRNA’s endogenous promoters. Each mirtron reached levels close to those of individual endogenous family members, and was able to largely rescue embryonic lethality of the corresponding miRNA-family deletion (to 85% and 75%, respectively) **(Fig. 2B, Fig. S2E-H)**, establishing the mirtron pathway as an efficient way to express Microprocessor-independent miRNAs that substitute for abundant, endogenous miRNAs.

**Fig. 2.**
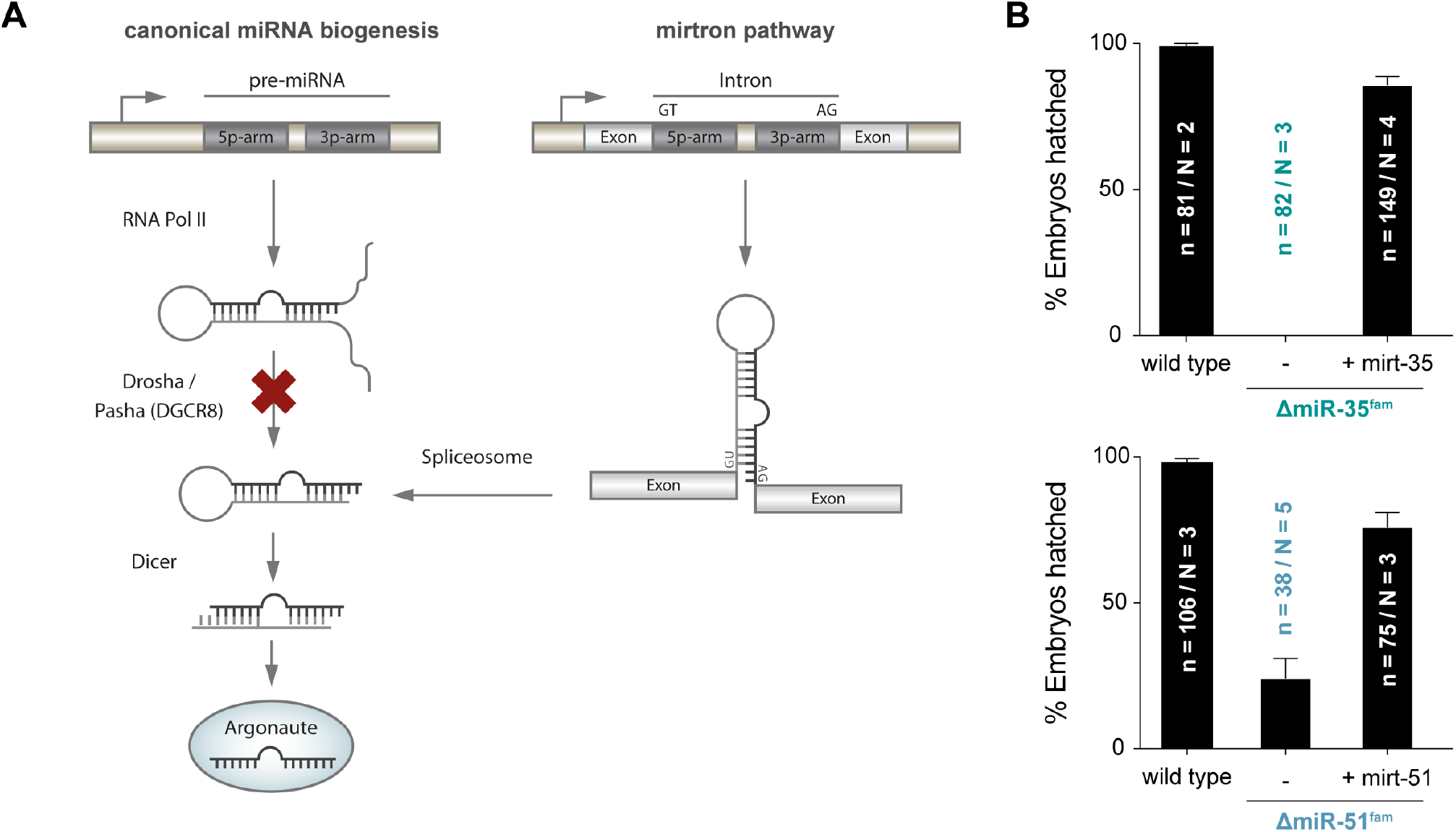
**A.** Schematic of the experimental strategy to test for miRNA sufficiency: a block in miRNA biogenesis at the Microprocessor level is bypassed by expression of miR-35 and miR-51 in form of mirtrons. **B.** Hatching rates of miR-35 and miR-51 family mutant embryos with or without a mirtron-version of a respective miRNA-family member (plotted as in 1A).

Simultaneous expression of mirt-35 and mirt-51 restored the ability of the RNAiD and *pash-1(ts)* Microprocessor-deficient backgrounds to produce morphologically-normal larvae in ~50% of cases **(Fig. 3)**; out of a theoretical maximum of ~63% given the imperfect rescue of each of the mirtron transgenes. The remaining embryos showed variable levels of rescue, with ~70% reaching later stages of development, albeit with morphogenesis defects **(Fig. S3A)**. Upon RNAiD treatment, both mirtrons were necessary to rescue embryogenesis. However, in the *pash-1(ts)* background, in which a higher fraction of maternal miR-35^fam^ and miR-51^fam^ members persists in early embryos **(Fig. S1G)**, each mirtron provided partial rescue on its own **(Fig. S3B)**.

**Fig. 3.**
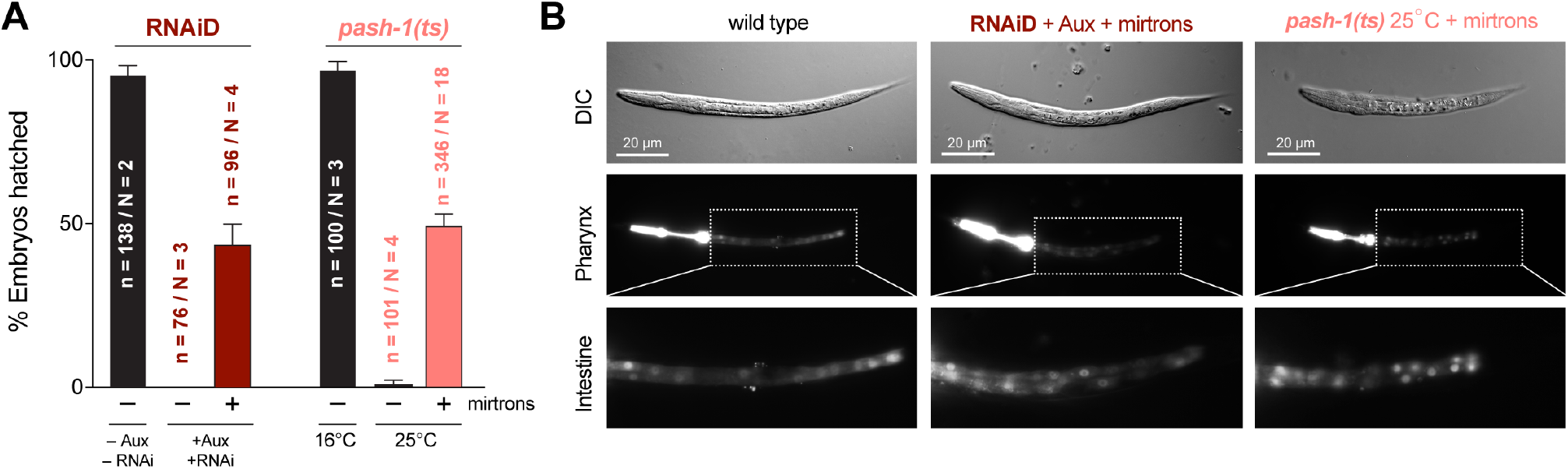
**A.** Hatching rates of the two independent Microprocessor-depleted animals (RNAiD or *pash-1(ts)*) with or without mirtron-35 and mirtron-51 (plotted as in 1A). **B.** Representative images of mirtron-rescued L1 larvae. Shown are DIC/Nomarski images as well as fluorescently labelled pharynx (*myo-2^prom^::mCherry*) and intestine (*elt-2^prom^::NLS-dsRed*). For comparison to arrested embryos without mirtrons see Fig. 1A and S1D.

The mirtron-rescued larvae formed all major organs and appeared morphologically normal, although they were slightly shorter and wider **(Fig. 3B, Fig. S3C-E)**, likely due to the lack of the miR-58 family (*11*). They also crawled and fed, indicating that the production of these two miRNAs accounts for the majority of the embryogenesis defects observed in Microprocessor-deficient *C. elegans*. However, rescued larvae rarely molted, and all died. To test whether these larvae could resume development upon regaining ability to produce miRNAs, we shifted mirtron-rescued *pash-1(ts)* L1 larvae back to the permissive temperature. This enabled a quarter of the larvae to become fertile adults, suggesting that L1 larvae rescued by mirt-35 and mirt-51 during embryogenesis are competent to complete development **(Fig. S3F)**, and that additional miRNAs, including but not limited to *lin-4* and *let-7* (*17*, *18*), are required for larval progression.

We assessed the level of miRNA depletion in the rescued L1 larvae by quantitative small RNA sequencing. Mirtron-rescued L1 larvae of the *pash-1(ts)* strain had only trace amounts of most miRNAs **(Fig. 4A)**. Notably, two miRNAs persisted, the miR-51^fam^ member miR-52, which is partially resistant to Microprocessor ablation, and miR-62, an endogenous mirtron without obvious function **(*10*)**. RNAiD treatment, which was very efficient for depletion of maternal and zygotic miRNAs up to the end of gastrulation, was less effective for miRNA depletion in L1 larvae, likely because auxin levels in late embryos are insufficient to trigger efficient degradation. Nevertheless, from the combination of the RNAiD and the *pash-1(ts)* experiments, we conclude that miR-35^fam^ and miR-51^fam^ are sufficient for embryonic development in the absence of Drosha and Pasha/DGCR8 activities.

**Fig. 4.**
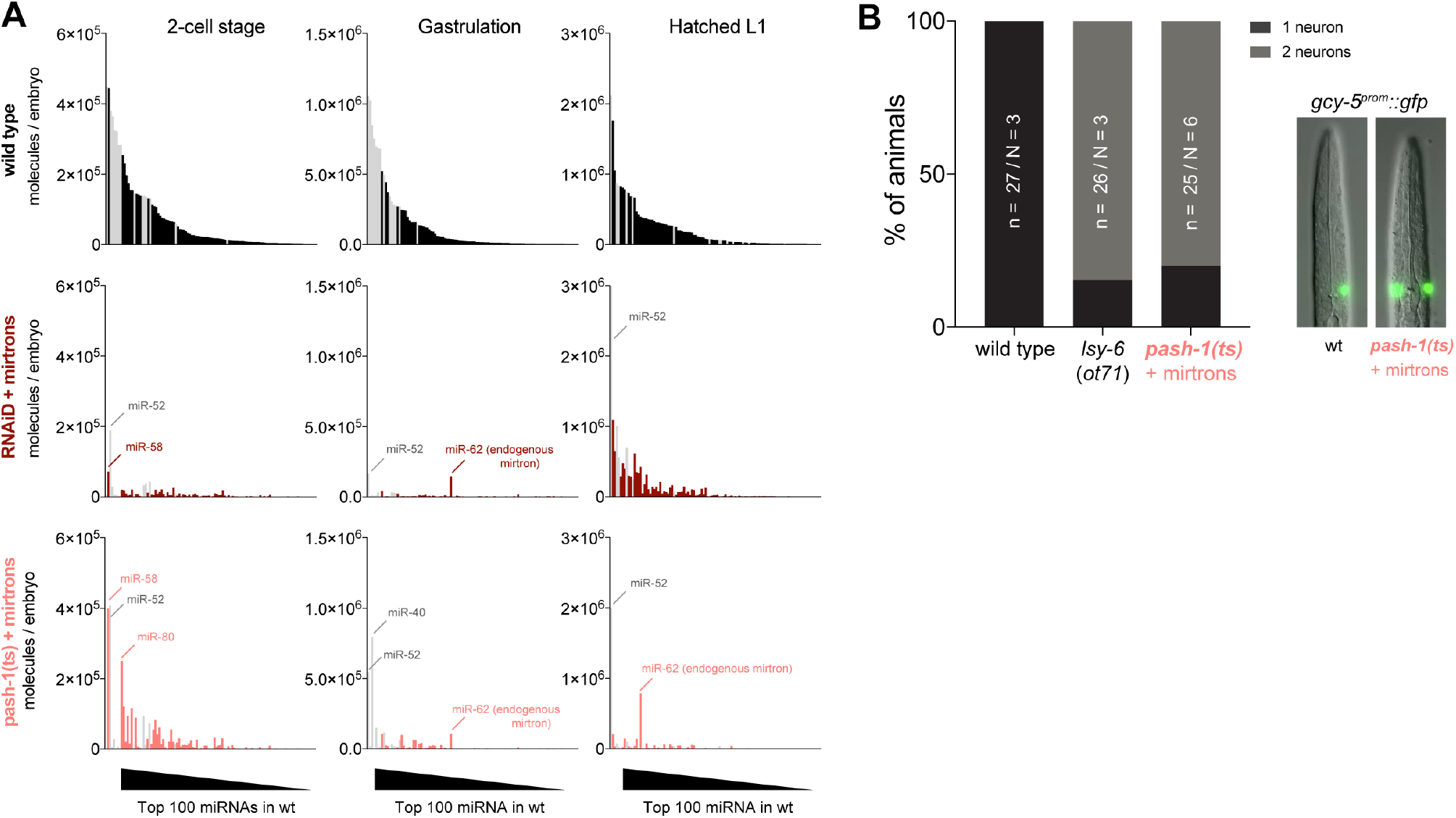
**A.** Absolute miRNA abundance in wt, RNAiD, and *pash-1(ts)* animals under restrictive conditions, at the indicated timepoints, measured by spike-in normalized small RNA-seq as in Fig. 1B. Shown are the top 100 miRNAs, ranked by abundance in wild-type samples. Remaining miRNAs of the miR-35 and -51 families are in gray. For comparison, data for 2-cell and gastrulation timepoints is replotted from Fig. 1B and S1G. **B.** Functional readout for the function of the late embryonic miRNA *lsy-6*. In wild-type animals only the right ASE neuron expresses *gcy-5:gfp*, but absence of functional *lsy-6* results in activation of the reporter in both the left and right neurons.

Mirtron-rescued larvae were still functionally depleted of miRNAs that act later in cell-type-specific manners. For instance, *lsy-6* is a miRNA necessary to lateralize the ASE pair of neurons (*19*). Mirtron-rescued *pash-1(ts)* larvae behaved identically to larvae bearing a genomic deletion of *lsy-6*, in their inability to properly specify the ASE lateral asymmetry **(Fig. 4B)**. Hence, these larvae, which lack practically all miRNAs except for the miR-35 and miR-51 families, provide an opportunity to explore the global contribution of such cell-type-specific miRNAs.

Both Drosha and Pasha/DGCR8 bind a variety of non-miRNA substrates (*20*–*22*) and perform regulatory functions through cleavage or binding of mRNAs (*23*–*26*). However, the biological significance of these functions remains unclear. As embryonic lethality caused by absence of Pasha and Drosha is rescued by addition of miR-35 and miR-51, we conclude that neither protein carries out essential miRNA-independent functions during *C. elegans* embryogenesis.

The miR-35^fam^ and -51^fam^ are broadly conserved, with miR-51 being a homolog of miR-100, the most ancient animal miRNA (*12*, *27*). Yet, the functions of these two miRNA families remain poorly understood and neither has been linked to mRNA targets that explain the penetrant embryonic lethality (*12*, *28*–*32*). Zebrafish miR-430 is expressed in the zygote, where it plays a role in maternal mRNA clearance (*4*). In contrast, miR-35^fam^ and -51^fam^ are present in the germline and, at least for miR-35^fam^, there is strong evidence that it is already active in the maternal germline (*31*, *32*). Moreover, the expression levels of the predicted targets of the miR-35^fam^ and -51^fam^ tend to stay constant during embryogenesis or even increase **(Fig. S4)** (*33*). These observations strongly suggest that miR-35^fam^ and -51^fam^ do not function in maternal mRNA clearance.

In summary, we showed that *C. elegans* embryos depleted of the Microprocessor arrest shortly after gastrulation, failing to undergo morphogenesis or to form organs. This arrest phenotype is earlier than that observed in Dicer-deficient zebrafish and closer to that of Pasha/DGCR8 mouse embryos, which die during gastrulation *(34)*. Two conserved miRNAs, miR-35^fam^ and -51^fam^, are sufficient to overcome this arrest in *C. elegans*. These miRNAs play a yet unidentified, likely conserved function in animal embryogenesis.

## Acknowledgments

We thank K. McJunkin, Cochella Lab members and G. Riddihough (Life Science Editors) for feedback on the manuscript, M. Asparuhova and K. Umundum for generation of transgenic animals, and Wormbase. Some strains were provided by the CGC (NIH grant P40 OD010440).

## Author contributions

PJD and LC designed the experiments, wrote the manuscript and prepared the figures, PJD performed wet-lab experiments, JW conducted bioinformatic analyses.

## Data and materials availability

Strains will be made available at CGC, sequencing data are deposited, the GEO accession number will be made available upon publication.

## Supplementary Material

### Materials and Methods

#### Animal maintenance

*Caenorhabditis elegans* was kept under standard conditions on NGM plates seeded with *Escherichia coli* OP50 at 20°C as previously described *(1)*, unless indicated otherwise. The Bristol strain N2 was used as a wild-type control. MiR-35 family mutant embryos were derived from self-fertilized, homozygous mutant mothers (nDf49, nDf50) that were maternally rescued. For analysis of the earliest miR-51 family mutant phenotypes, we sought to reduce the maternal load of miR-51 family miRNAs (as embryos can only be obtained from mothers expressing at least one mir-51 family member). To this end, mutant embryos were derived from mothers expressing miR-54-56 as well as an early embryonic *mir-35p::GFP* reporter from an extrachromosomal array, while all genomic copies of miR-51 family miRNAs were deleted (nDf58, nDf67, n4100). These arrays tend to be lowly expressed or silenced in the *C. elegans* germline, and are only segregated to a fraction of the offspring. Therefore, miR-51 family mutant embryos obtained from these mothers upon self-fertilization were selected for lack of the rescuing array by absence of *mir-35p::GFP* expression, and subsequently analyzed. A full list of strains used and generated in this study is provided in Table S1.

#### Mirtron design and expression

Production of a miRNA of interest via the mirtron pathway involves the challenge of designing an intron that resembles a pre-miRNA hairpin and simultaneously allows for efficient splicing. To generate mirtron-versions of miR-35 or miR-51, the 6 nucleotides at the 3’ end of the respective pre-miRNA-hairpin were modified to create a 3’ splice site and match the consensus sequence of 12 annotated endogenous *C. elegans* mirtrons **(Fig. S2A)** *(2)*. Because intron borders correspond to the ends of mature miRNAs, the 5’ arm of mirtrons invariably starts with a G, which negatively impacts loading into Argonaute *(3)*. Thus, mature mirtrons are preferably incorporated into the 3’ arm. As interaction of miRNAs with cognate target mRNAs is largely determined by the seed region (nts 2-8), minor alterations in the 3’ end of a miRNA are expected to have little impact on its target repertoire. Upon introduction of a functional 3’ splice site into the 3p-arm of the pre-miRNA, compensatory changes were introduced in the 5p arm to retain hairpin secondary structure. This was guided by secondary structure predictions using the RNAfold Two-State-Folding algorithm for RNA at 20°C *(4),* yielding a mirtron-hairpin with a secondary structure highly similar to its endogenous pre-miRNA-hairpin equivalent **(Fig. S2C)**, the resulting sequence was used to replace the middle intron of GFP in the commonly employed *C. elegans* vector pPD75.95 (Fire lab). To recapitulate the spatio-temporal expression pattern of endogenous miRNAs, the promoter sequence of the respective endogenous miRNA was used to drive mirtron expression (2.0 kb upstream of pre-mir-35 or 2.5kb upstream of pre-mir-52, respectively). Conveniently, GFP-fluorescence serves as an internal control for correct splicing of the inserted mirtron. Functionality of mirtrons was assayed by testing their ability to rescue an otherwise lethal deletion of the respective miRNA family. For an estimate of mirtron-expression levels relative to endogenous miRNA family members, TaqMan (ThermoFisher) RT-qPCR was employed using RT-primers as well as probes specific to the 3’ end variations in mirtrons and endogenous miRNA counterparts. Expression levels were calculated using the ΔΔCq method relative to U18 snoRNA as a reference transcript.

#### Genome engineering

To generate AID-tagged copies of *pash-1* and *drsh-1,* animals were subjected to Cas-9 mediated genome engineering via RNP-microinjection following the Co-CRISPR approach described by *(5)*, with modifications from *(6)*. Briefly, the Alt-R CRISPR/Cas9 system (Integrated DNA Technologies) was employed and animals were injected with a mixture containing 300mM KCl, 20mM HEPES, 4μg/μl recombinant Cas9 (from *S. aureus,* purified in-house), 500ng/μl TracerRNA, 100ng/μl crRNA targeting the Co-CRISPR marker gene dpy-10, and 15ng/μl dpy-10->rol-6 repair template oligo. Additionally, 100 ng/μl crRNA against the locus of interest as well as 100-200ng/μl dsDNA repair template encoding the desired modification with around 50bp flanking homology arms were added to the mix. Note that for efficient degradation of DRSH-1 we needed to insert AID degrees at the N- and C-termini simultaneously as either degree alone was insufficient for depletion. Guide RNAs used and sequences inserted were as follows:

*drsh-1* – N-terminal AID-tag – guide sequence used = 5’- TTAGATTTTCATTTAGATGT-3’ – locus post-edit:

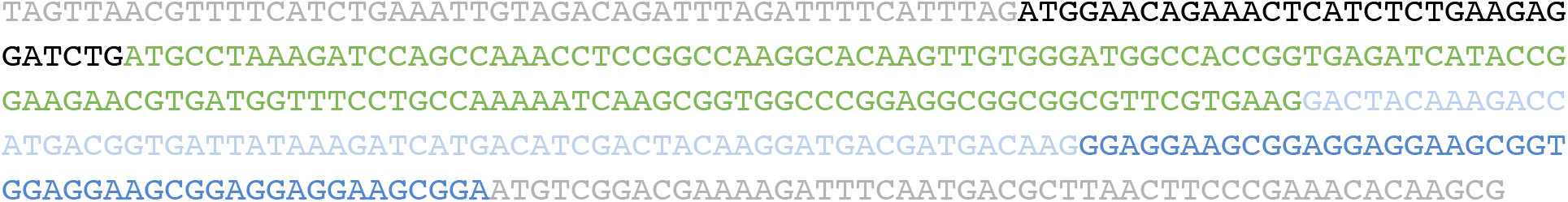

*drsh-1* – C-terminal AID-tag – guide sequence used = 5’-GATACCAGCGACTAATTACG-3’ – locus post-edit:

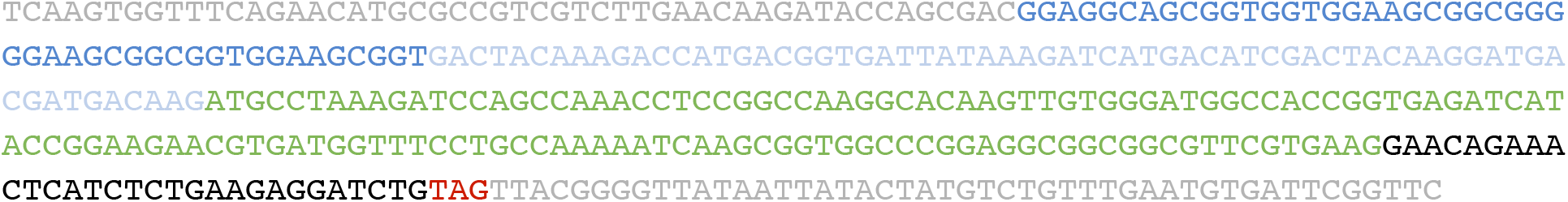

*pash-1* – C-terminal AID-tag – guide sequence used = 5’-AGGTGAATATACTATTTGTG-3’ – locus post-edit:

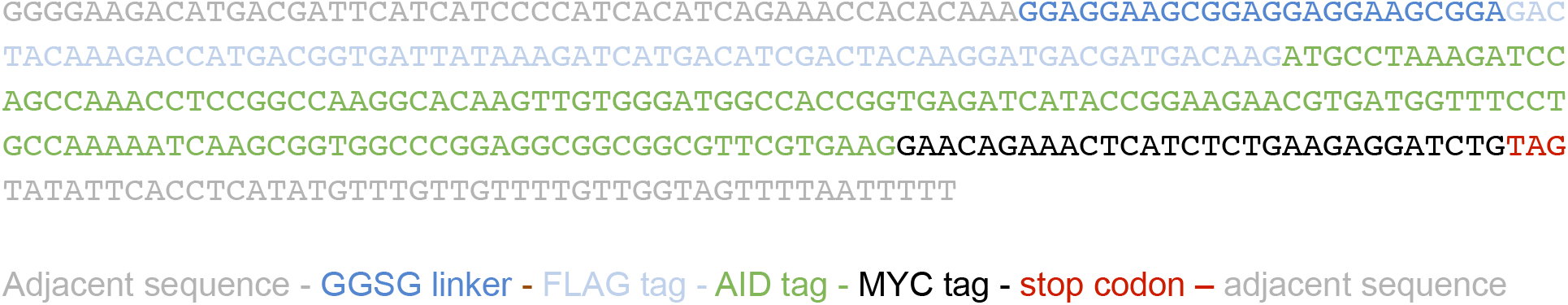

#### Pash-1(ts) experiments

Animals bearing the *pash-1(ts)* allele *mj100* were kept constantly at the permissive temperature of 16°C. *pash-1(ts)* animals expressing the mirtron-51 transgene were selected every few generations for high levels of the co-expression marker *elt-2p::dsRed*, as this transgene was prone to undergo silencing. For experiments, L4 stage larvae were transferred to a fresh plate and shifted to 25°C approximately 16h later. In order to achieve maximal depletion of maternal miRNAs, mothers were kept at the restrictive temperature an additional 7h before harvesting early embryos (1-4 cell stage). Extending the time at the restrictive temperature was prohibited by the rapidly deteriorating health of gravid *pash-1(ts)* adults expressing mirtrons at 25°C. For backshift-experiments with mirtron-rescued *pash-1(ts)* animals, hatched L1s that developed under restrictive conditions were transferred to a fresh plate within 1h after hatching, allowed to develop at 16°C, and scored for their ability to reach adulthood within 7 days.

#### RNAiD experiments

Animals were kept constantly on standard NGM plates seeded with *E. coli* OP50 at 20°C. For combined AID and RNAi treatment, NGM plates were supplemented with 1mM Carbenicillin, 1mM IPTG and 4mM Auxin (3-Indoleacetic acid, Sigma #I2886). RNAiD plates were seeded with *E.coli* HT115 expressing dsRNA which elicits an RNAi response against *pash-1* mRNA (clone from the ORFeome-RNAi v1.1 (Vidal) library, ORF-ID T22A3.5). To ensure efficient mRNA knockdown, RNAi bacteria were grown freshly for every experiment in liquid LB culture, the medium was brought to 1mM IPTG around 1h before pelleting bacteria, and plates were seeded with a 20X concentrate (as Auxin inhibits bacterial growth). For experiments, L4s were transferred to RNAiD plates, embryos were harvested 24h later, and assayed as described below. While the AID-system rapidly depletes proteins within minutes to hours *(7)*, extending the Auxin treatment to 24h was required to ensure near-complete elimination of maternal miRNAs. Surprisingly, overall health of mirtron-expressing gravid adults upon RNAiD treatment not adversely affected, compared to *pash-1(ts)* animals, eventually owed to the temperature difference and/or deleterious side-effects of PASH-1(TS) at 25°C. Efficiency of RNAi-mediated *pash-1* mRNA knockdown was assessed via RT-qPCR following the procedure described in *(8)*. Briefly, 5-10 embryos in the 2-cell stage were transferred to about 1 μl of Lysis Buffer (5mM Tris-HCl pH 8.0, 0.25 mM EDTA and 1 mg/mL Proteinase K, 0.5% Triton X-100, 0.5% Tween20) using a thin glass needle. These samples were subjected to 10 min digestion at 65°C before heat inactivation of proteinase K for 1 min at 85°C. Next, crude lysates were reverse transcribed before performing qPCR using the GoTAQ qPCR Mastermix (Promega) according to the manufacturer’s instructions. Relative expression was calculated according to the ΔΔCq-method using *cdc-42* as a reference gene, the following primer sequences were employed:

*pash-1*-F: 5’-GCCTTCGAGAAAACGGGGAA-3’; *pash-1*-R: 5’-TGGCTCCCATTTCGGAGATT-3’;
*drsh-1*-F: 5’-TGAGCTGGCTTTGGCTAATCT-3’; *drsh-1*-R: 5’-ACCCCGTAATTAGTCGCTGG-3’
*cdc-42*-F: 5’-TGGGTGCCTGAAATTTCGC-3’; *cdc-42*-R: 5’-CTTCTCCTGTTGTGGTGGG-3’

#### Western Blotting

To assess the extent of protein degradation of PASH-1:AID:MYC and DRSH-1:AID:MYC upon Auxin treatment, L4s were transferred to Auxin plates prepared as described above. 24h later, gravid adults were collected in Lämmli-buffer containing 2.5% ß-Mercaptoethanol (v/v), and subjected to multiple cycles of snap-freezing followed by boiling until embryos were disrupted. Proteins were separated via SDS-PAGE and transferred onto a Nitrocellulose membrane. As primary antibodies either monoclonal mouse anti-myc (clone 46A, Merck Millipore, catalog number 05-724, diluted 1:2000) or a polyclonal rabbit anti-gamma tubulin (Abcam, catalog number ab50721, diluted 1:1000) were used. As secondary antibody, anti-mouse IgG HRP-linked Antibody (Cell Signaling Technology, #7076, diluted 1:2,000) or anti-rabbit IgG HRP-linked Antibody (Cell Signaling Technology, #7074, dilution of 1:2000) was applied followed by visualization using ECL reagent (Thermo Scientific).

#### Hatching assays

Embryos were obtained from day 1 gravid adults reared as described above, by slicing mothers in a drop of M9 buffer on a microscopic slide. 2-cell embryos of normal size were transferred to standard NGM plates or RNAiD plates and allowed to develop at the respective temperatures. Hatched animals were collected for subsequent analysis (backshift experiments, microscopy, or small RNA sequencing) from plates within 1h after hatching. The final hatching rate was assessed >24h after embryo collection by scoring the fraction of animals that successfully escaped the eggshell. To obtain samples at specific developmental stages for small RNA sequencing, 2-cell embryos were harvested via slicing, collected or allowed to developed on NGM plates before being selected manually by stage at given time points (Gastrulation: 4-5h after 2-cell stage).

#### Microscopy

For phenotypic analysis of embryonic arrest or hatched L1 larvae, animals were mounted on a thin agar-pad on a microscopic slide sealed with a coverslip. Images were recorded at 400x magnification using an AxioImager Z2 (Zeiss) equipped with DIC and fluorescence optics, and analyzed via ImageJ. Arrest phenotypes were scored as follows: cell mass = embryo fails to initiate bean stage; morphogenesis = embryo begins to elongate but fails to reach 2-fold stage; elongation = embryo completes 2-fold stage but arrests before hatching. Body measurements are based on DIC micrographs. Body length was measured as the length of a segmented path running through the mid body axis from head to tail. Body width was assessed as the arithmetic mean of 3 independent width measurements in the anterior, middle, and posterior part of each animal. Pharynx length was determined by the length of a segmented path running through the pharynx middle axis in animals expressing *myo-2^prom^::mCherry*. Mean and range are plotted for each genotype; unpaired t-test was used for statistical comparison.

#### Small RNA sequencing

To profile miRNA levels, a modified version of the small RNA sequencing protocol described in (*9*) was performed. Samples containing each between 25 and 100 embryos (yielding about 5 to 20 ng total RNA) were collected at indicated stages and snap-frozen in liquid nitrogen. A series of eight RNA spike-In oligos spanning a 500-fold range of concentrations *(10)* were added on a per embryo basis, and total RNA was extracted using TRIzol Reagent (Invitrogen) according to the recommendations by the manufacturer for cell samples. To ensure complete disruption of embryos, samples were snap-frozen and thawed multiple times before proceeding with the extraction. After total RNA purification samples were treated as described in (*9*), with two important changes: A) amounts of 3’ linker and 5’ linker were reduced to a final concentration of 500 nM to reduce undesired amplification products. B) barcodes and random nucleotides serving as unique molecular identifiers were introduced into the 3’ linker to circumvent low-input related contamination and over-amplification issues, resulting in ligation products of the following sequence:

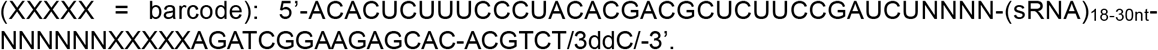

After sufficient PCR-amplification as observed by SYBR green derived qPCR signal (requiring around 16-20 cycles), libraries were size-selected on an agarose gel to remove adaptor dimers and sequenced on a HiSeqV4 platform (Illumina). Small RNA reads were mapped and processed as in (*9*). Modifications include addition of 3’adaptor-barcode demultiplexing, which was added after trimming adaptor sequence in the Nextflow-based pipeline (Tommaso et al., 2017). The developed Nextflow pipeline can be found at: https://github.com/lengfei5/smallRNA_nf/tree/master/dev_sRBC. To accurately quantify sRNAs, both mapped sequences and associated random nucleotides (functioning as UMIs) within the raw reads were quantified for miRNAs and piRNAs as well as spike-ins. To alleviate PCR amplification bias, UMI counts were used and normalized to spike-in as described in *(10)*, allowing for absolute quantification of miRNA content per embryo even under conditions in which most miRNAs are depleted. In addition, as piRNAs are not affected across conditions and can thus be considered an internal control, data was normalized in parallel using total number of piRNA reads, yielding results highly similar to the ones obtained by spike-in normalization and providing additional confidence in the quantitative nature of the data. Scripts for the analysis can be found at: https://github.com/lengfei5/smallRNA_analysis_philipp/tree/master/scripts. Every round of library preparation included one sample of a comparable amount of *Arabidopsis thaliana* total RNA to assess the extent of potential contamination. The average counts for reads mapping to *C. elegans* miRNAs in contamination control samples (~2% of the wild-type miRNA content) across four independent library preparations were subtracted from all samples as a background correction. For background-corrected, spike-in normalized, miRNA counts in molecules per embryo, see supplemental table S2. Raw data from sRNAseq experiments has been deposited in the NCBI-GEO database under the accession number: GSE153233.

## Supplementary Figures

**Figure S1.**
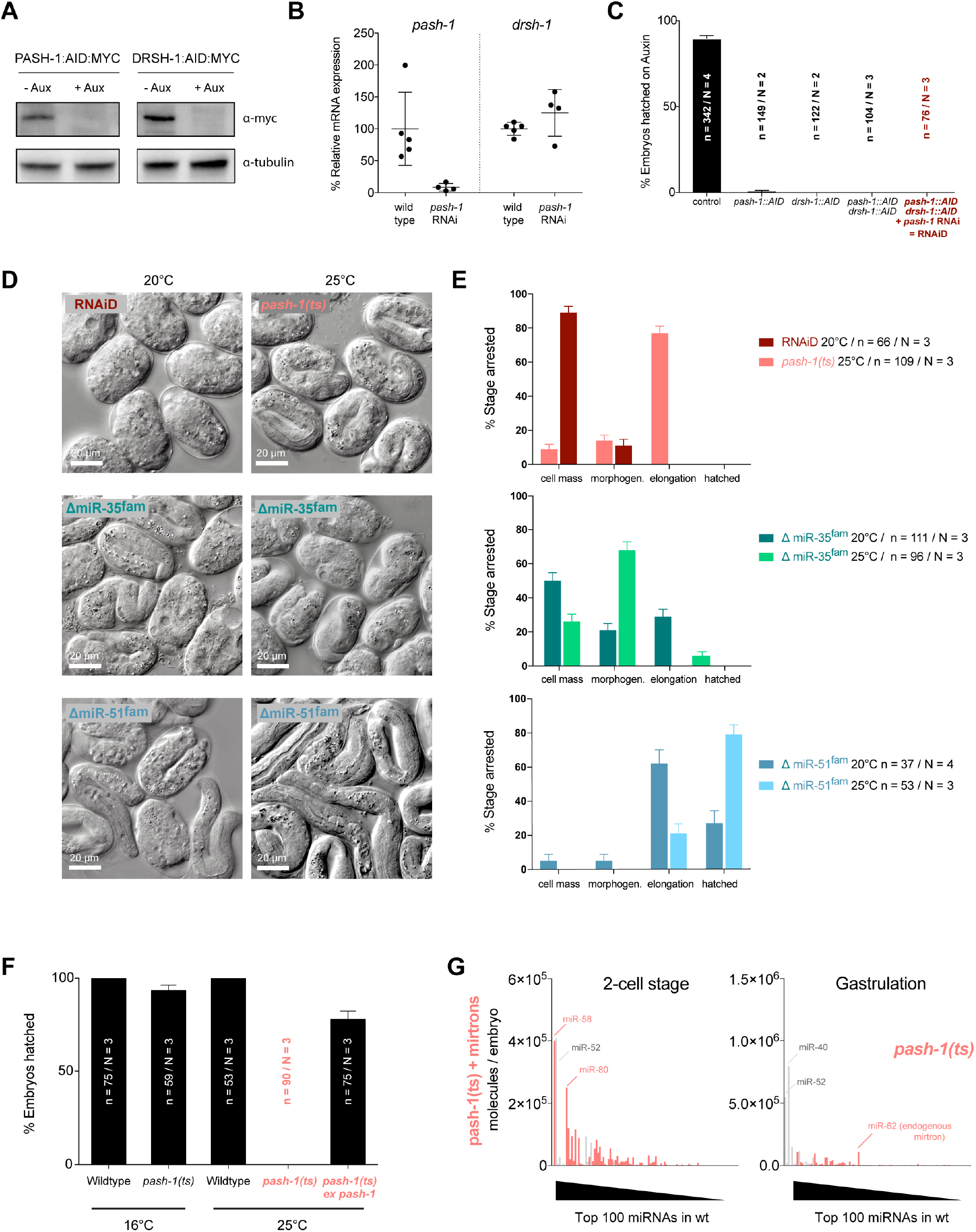
**A**) Western Blot against endogenous, AID-Myc-tagged PASH-1 and DRSH-1 in gravid adults, 24h post Auxin treatment. **B**) mRNA levels measured by RT-qPCR in 2-cell embryos, 24h post *pash-1* RNAi treatment compared to wild type under standard conditions. **C**) Hatching assay with various strains bearing AID-tagged Microprocessor components upon Auxin-treatment. For comparison, data for control and RNAiD conditions are replotted from Fig. 1A. **D, E**) Images and quantification of arrest phenotype of RNAiD, *pash-1(ts)*, miR-35^fam^, and miR-51^fam^ mutant embryos at 20°C and 25°C. **F**) Hatching assay of embryos carrying the *pash-1(ts)* temperature-sensitive allele *mj100*. Penetrant embryonic lethality at the restrictive temperature 25°C is rescued by expression of wild-type *pash-1* from an extra-chromosomal array. **G**) miRNA-profile of *pash-1(ts)* embryos at the 2-cell stage and at the end of gastrulation as determined by small RNA sequencing using spike-in oligos for quantitative measure of absolute miRNA levels across samples. The top 100 most abundant miRNAs are shown, ranked by abundance in wild-type embryos. Remaining miRNAs of the miR-35^fam^ and miR-51^fam^ are in gray.

**Figure S2.**
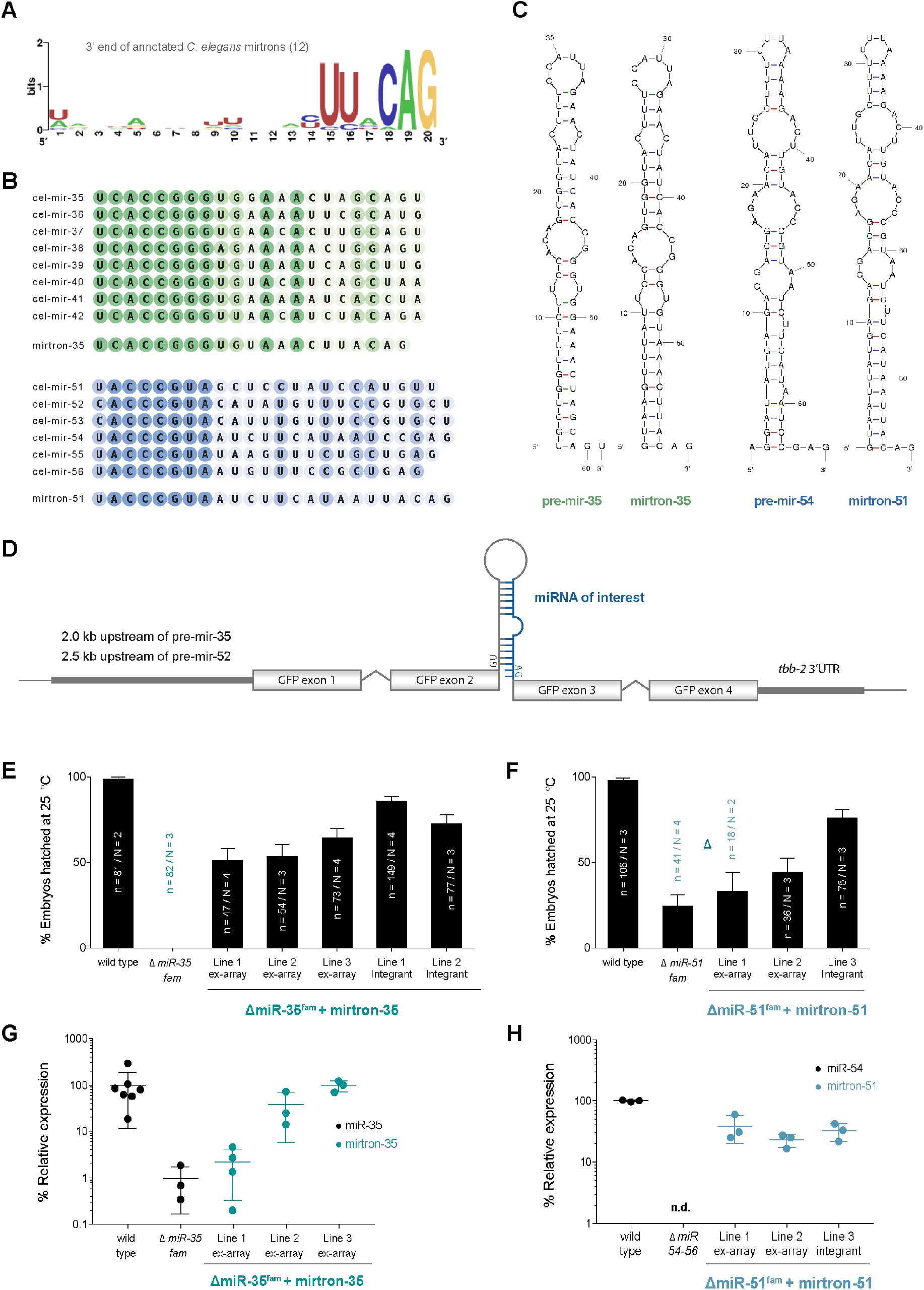
**A**) Consensus sequence at the 3’ end of all 12 annotated *C. elegans* mirtrons *(2)*. **B**) Sequence of miR-35 and miR-51 family miRNAs and the corresponding designed mirtrons. **C**) Pre-miRNA hairpin structure for endogenous miRNA and mirtron counterpart as predicted by the UnaFold TwoState folding algorithm for RNA at 20°C *(4)*; the base for mirtron-51 was the family member miR-54 as it required fewer changes to be turned into a mirtron. **D**) Schematic design of mirtron-expressing transgenes. The endogenous promoter sequence upstream of pre-mir-35 (2.0 kb) or pre-mir-52 (2.5kb) drives expression of a GFP transcript bearing the mirtron in its middle intron, followed by an inert *tbb-2* (tubulin) 3’ UTR. **E, F**) Hatching rates of embryos bearing deletions for the miR-35 family (nDf49, nDf50), or the miR-51 family (nDf58, nDf67, n4100), respectively. Shown are multiple independent transgenic lines expressing the respective mirtron from an extra-chromosomal array or the integrated transgenes that were used for subsequent experiments. **G, H**) Expression of mirtron-35 or -51 and the respective endogenous counterparts (miR-35 or -54) as measured by TaqMan RT-qPCR in individual embryos at the comma stage.

**Figure S3.**
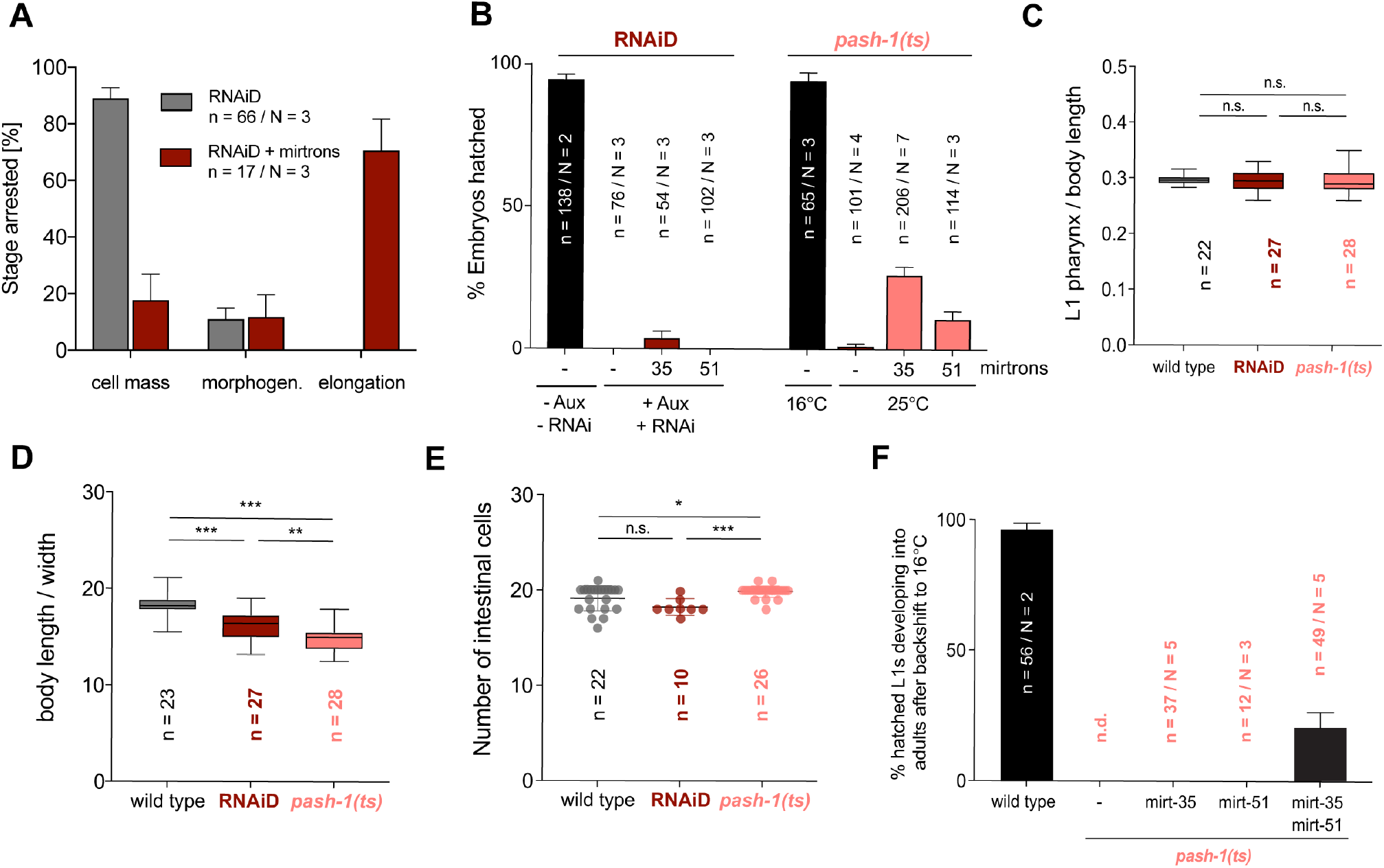
**A**) Stage of arrest in the ~50% of RNAiD embryos with mirtron-35 and -51 that fail to hatch, compared to embryos without mirtrons. **B**) Hatching assay of microprocessor-deficient animals expressing either mirtron-35 or mirtron-51 individually. **C**-**E**) Body measurements of mirtron-rescued RNAiD or *pash-1(ts)* animals compared to wild type under standard conditions. Data is derived from micrographs using ImageJ. Pharynx length was measured using the expression area of a *myo-2^prom^::mCherry* reporter. Number of intestinal cells was scored by expression of a nuclear *elt-2^prom^::dsRed* reporter. Mean and range are plotted for each genotype; statistical significance was determined by an unpaired t-test, significance levels are: P>0.05 = n.s.; P<0.05 = ✽; P<0.01 = ✽✽; P<0.001 = ✽✽✽. **F**) *pash-1(ts)* backshift experiment: miRNA-depleted embryos that developed at the restrictive temperature of 25°C and expressed mirt-35/51 individually or in combination, were transferred to the permissive temperature 16°C directly after hatching. Survival to adulthood was scored within the next 7 days.

**Figure S4.**
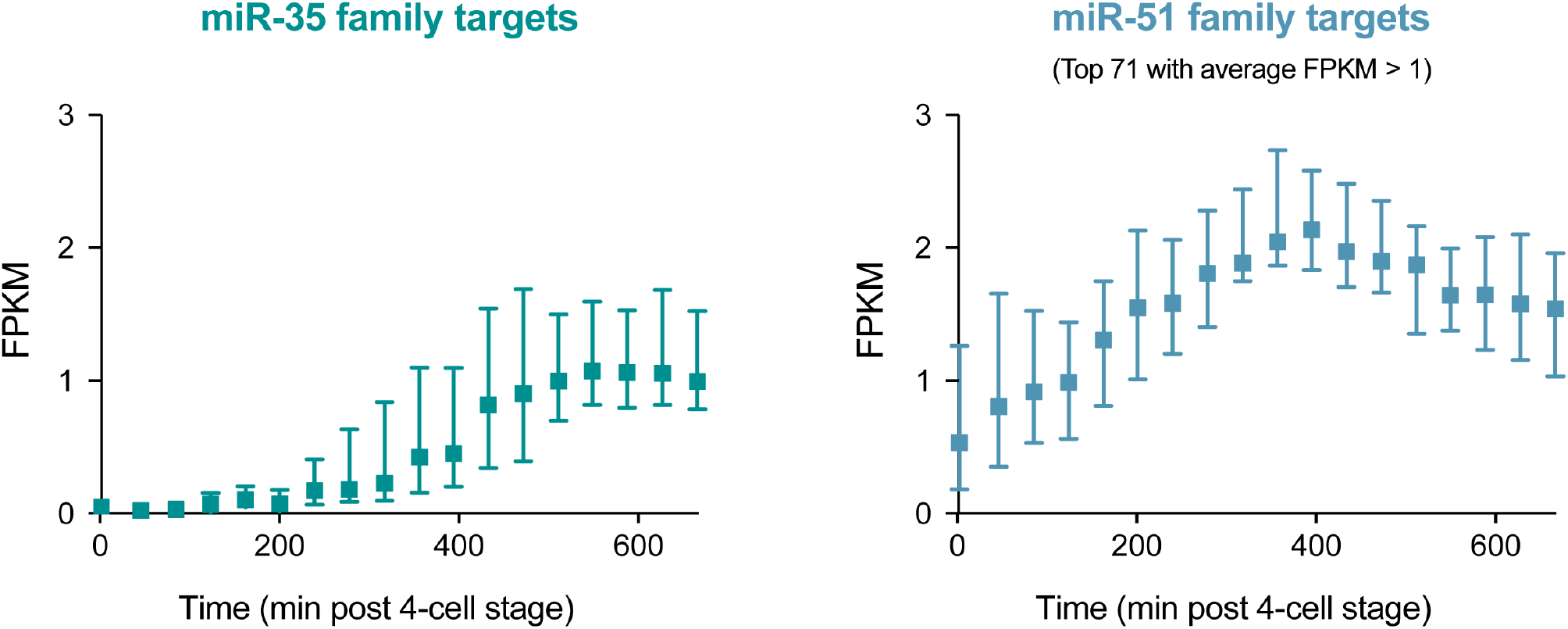
Embryonic expression time course of transcripts bearing conserved miR-35/51 family miRNA binding sites as predicted by TargetScan Worm 6.2 *(11, 12)*. Shown are median expression and 95% confidence intervals. For miR-35 family all 89 predicted targets are shown. For miR-51 family targets the top 71 expressing transcripts (out of 264 called targets in the dataset) are shown, selected based on an average FPKM value > 1 across all timepoints. The most abundant predicted miR-51^fam^ target *rps-20* (ribosomal protein, small subunit) has been removed as an outlier due to extreme abundance. Data derived from *(13)*.

## References

1. D. P. Bartel, Metazoan MicroRNAs. Cell. 173, 20–51 (2018).

2. C. Alberti, L. Cochella, A framework for understanding the roles of miRNAs in animal development. Development. 144, 2548–2559 (2017).

3. A. J. Giraldez et al., MicroRNAs regulate brain morphogenesis in zebrafish. Science. 308, 833–838 (2005).

4. A. J. Giraldez et al., Zebrafish MiR-430 promotes deadenylation and clearance of maternal mRNAs. Science. 312, 75–79 (2006).

5. B. Fromm et al., MirGeneDB 2.0: the metazoan microRNA complement. Nucleic Acids Research. 48, D132–D141 (2019).

6. A. M. Denli, B. B. J. Tops, R. H. A. Plasterk, R. F. Ketting, G. J. Hannon, Processing of primary microRNAs by the Microprocessor complex. Nature, 5 (2004).

7. C. Rios, D. Warren, B. Olson, A. L. Abbott, Functional analysis of microRNA pathway genes in the somatic gonad and germ cells during ovulation in C. elegans. Developmental Biology. 426, 115–125 (2017).

8. L. Zhang, J. D. Ward, Z. Cheng, A. F. Dernburg, The auxin-inducible degradation (AID) system enables versatile conditional protein depletion in C. elegans. Development. 142, 4374–4384 (2015).

9. N. J. Lehrbach et al., Post-developmental microRNA expression is required for normal physiology, and regulates aging in parallel to insulin/IGF-1 signaling in C. elegans. RNA. 18, 2220–2235 (2012).

10. E. A. Miska et al., Most Caenorhabditis elegans microRNAs Are Individually Not Essential for Development or Viability. PLoS Genet. 3, e215 (2007).

11. E. Alvarez-Saavedra, H. R. Horvitz, Many families of C. elegans microRNAs are not essential for development or viability. Curr. Biol. 20, 367–373 (2010).

12. W. R. Shaw, J. Armisen, N. J. Lehrbach, E. A. Miska, The Conserved miR-51 microRNA Family Is Redundantly Required for Embryonic Development and Pharynx Attachment in Caenorhabditis elegans. Genetics. 185, 897–905 (2010).

13. C. Alberti et al., Cell-type specific sequencing of microRNAs from complex animal tissues. Nat Meth. 15, 283–289 (2018).

14. N. J. Martinez et al., Genome-scale spatiotemporal analysis of Caenorhabditis elegans microRNA promoter activity. Genome Research. 18, 2005–2015 (2008).

15. K. Okamura, J. W. Hagen, H. Duan, D. M. Tyler, E. C. Lai, The mirtron pathway generates microRNA-class regulatory RNAs in Drosophila. Cell. 130, 89–100 (2007).

16. J. G. Ruby, C. H. Jan, D. P. Bartel, Intronic microRNA precursors that bypass Drosha processing. Nature. 448, 83–86 (2007).

17. R. C. Lee, R. L. Feinbaum, V. Ambros, The C. elegans heterochronic gene lin-4 encodes small RNAs with antisense complementarity to lin-14. Cell. 75, 843–854 (1993).

18. B. J. Reinhart et al., The 21-nucleotide let-7 RNA regulates developmental timing in Caenorhabditis elegans. Nature. 403, 901–906 (2000).

19. R. J. Johnston Jr, O. Hobert, A microRNA controlling left/right neuronal asymmetry in Caenorhabditis elegans. Nature. 426, 845–849 (2003).

20. S. Macias et al., DGCR8 HITS-CLIP reveals novel functions for the Microprocessor. Nat. Struct. Mol. Biol. 19, 760–766 (2012).

21. B. Kim, K. Jeong, V. N. Kim, Genome-wide Mapping of DROSHA Cleavage Sites on Primary MicroRNAs and Noncanonical Substrates. Molecular Cell. 66, 258–269.e5 (2017).

22. N. Gromak et al., Drosha Regulates Gene Expression Independently of RNA Cleavage Function. Cell Reports. 5, 1499–1510 (2013).

23. J. Han et al., Posttranscriptional Crossregulation between Drosha and DGCR8. Cell. 136, 75–84 (2009).

24. R. Triboulet, H.-M. Chang, R. J. LaPierre, R. I. Gregory, Post-transcriptional control of DGCR8 expression by the Microprocessor. RNA. 15, 1005–1011 (2009).

25. S. Kadener et al., Genome-wide identification of targets of the drosha-pasha/DGCR8 complex. RNA. 15, 537–545 (2009).

26. P. Knuckles et al., Drosha regulates neurogenesis by controlling Neurogenin 2 expression independent of microRNAs. Nat Neurosci. 15, 962–969 (2012).

27. A. Grimson et al., Early origins and evolution of microRNAs and Piwi-interacting RNAs in animals. Nature. 455, 1193–1197 (2008).

28. K. McJunkin, V. Ambros, The embryonic mir-35 family of microRNAs promotes multiple aspects of fecundity in Caenorhabditis elegans. G3 Genes|Genomes|Genetics. 4, 1747–1754 (2014).

29. K. McJunkin, V. Ambros, A microRNA family exerts maternal control on sex determination in C. elegans. Genes & Development. 31, 422–437 (2017).

30. R. Sherrard et al., miRNAs cooperate in apoptosis regulation during C. elegans development. Genes & Development. 31, 209–222 (2017).

31. A. T. Tran et al., MiR-35 buffers apoptosis thresholds in the C. elegans germline by antagonizing both MAPK and core apoptosis pathways. Cell Death and Differentiation. 26, 2637–2651 (2019).

32. M. A. Doll, N. Soltanmohammadi, B. Schumacher, ALG-2/AGO-Dependent mir-35Family Regulates DNA Damage-Induced Apoptosis Through MPK-1/ERK MAPK Signaling Downstream of the Core Apoptotic Machinery in Caenorhabditis elegans. Genetics. 213, genetics. 302458.2019–194 (2019).

33. M. E. Boeck et al., The time-resolved transcriptome of C. elegans. Genome Research. 26, 1441–1450 (2016).

34. Y. Wang, R. Medvid, C. Melton, R. Jaenisch, R. Blelloch, DGCR8 is essential for microRNA biogenesis and silencing of embryonic stem cell self-renewal. Nature Genetics. 39, 380–385 (2007).

## Supplementary References

1. S. Brenner, The genetics of Caenorhabditis elegans. Genetics. 77, 71–94 (1974).

2. W. J. Chung et al., Computational and experimental identification of mirtrons in Drosophila melanogaster and Caenorhabditis elegans. Genome Research. 21, 286–300 (2011).

3. F. Frank, N. Sonenberg, B. Nagar, Structural basis for 5'-nucleotide base-specific recognition of guide RNA by human AGO2. Nature. 465, 818–822 (2010).

4. N. R. Markham, M. Zuker, DINAMelt web server for nucleic acid melting prediction. Nucleic Acids Research. 33, W577–81 (2005).

5. A. Paix, A. Folkmann, D. Rasoloson, G. Seydoux, High Efficiency, Homology-Directed Genome Editing in Caenorhabditis elegans Using CRISPR-Cas9 Ribonucleoprotein Complexes. Genetics. 201, 47–54 (2015).

6. H. Prior, A. K. Jawad, L. MacConnachie, A. A. Beg, Highly Efficient, Rapid and Co-CRISPR-Independent Genome Editing in Caenorhabditis elegans. G3: Genes|Genomes|Genetics. 7, 3693–3698 (2017).

7. L. Zhang, J. D. Ward, Z. Cheng, A. F. Dernburg, The auxin-inducible degradation (AID) system enables versatile conditional protein depletion in C. elegans. Development. 142, 4374–4384 (2015).

8. K. Ly, S. J. Reid, R. G. Snell, Rapid RNA analysis of individual Caenorhabditis elegans. MethodsX. 2, 59–63 (2015).

9. C. Alberti et al., Cell-type specific sequencing of microRNAs from complex animal tissues. Nat Meth. 15, 283–289 (2018).

10. S. Lutzmayer, B. Enugutti, M. D. Nodine, Novel small RNA spike-in oligonucleotides enable absolute normalization of small RNA-Seq data. Sci Rep. 7, 5913 (2017).

11. B. P. Lewis, C. B. Burge, D. P. Bartel, Conserved seed pairing, often flanked by adenosines, indicates that thousands of human genes are microRNA targets. Cell. 120, 15–20 (2005).

12. C. H. Jan, R. C. Friedman, J. G. Ruby, D. P. Bartel, Formation, regulation and evolution of Caenorhabditis elegans 3'UTRs. Nature. 469, 97–101 (2011).

13. M. E. Boeck et al., The time-resolved transcriptome of C. elegans. Genome Research. 26, 1441–1450 (2016).

